# 3D-Aligner: An advanced computational tool designed to correct image distortion in expansion microscopy for precise 3D reconstitution and quantitative analysis

**DOI:** 10.1101/2024.08.28.610199

**Authors:** Jonathan Loi, Dhaval Ghone, Xiaofei Qu, Aussie Suzuki

**Author notes:** **Corresponding Author** Aussie Suzuki, University of Wisconsin-Madison.

## Abstract

Expansion Microscopy (ExM) is an innovative and cost-effective super-resolution microscopy technique that has become popular in cell biology research. It achieves super-resolution by physically expanding specimens. Since its introduction, ExM has undergone continuous methodological developments to enhance its resolution and labeling capabilities. However, ExM imaging often encounters sample drift during image acquisition due to the physical movement of the expanded hydrogel, posing a significant challenge for accurate image reconstruction. Despite many proposed experimental solutions to mitigate sample drift, a universal solution has yet to be established. In response to this challenge, we developed 3D-Aligner, an advanced and user-friendly image analysis tool designed to computationally correct drift in ExM images for precise three-dimensional image reconstruction and downstream quantification. We demonstrate that 3D-Aligner effectively determines and corrects drift in ExM images with different expansion rates and various fluorescently labeled biological targets, showcasing its capabilities and robustness in drift correction. Additionally, we validate the precision of 3D-Aligner by comparing drift values across different labeled targets and highlight the importance of drift correction in quantification of biological structures.

## Introduction

Fluorescence microscopy in biological research has become an essential tool due to its contributions towards myriad biological and biomedical discoveries. Its high sensitivity and specificity for labeling biological targets have empowered researchers to investigate the subcellular locations, structures, and functions of these targets within cells and tissues. Despite widespread use, fluorescence microscopy has faced a fundamental limit to its optical resolution to ∼250 nm laterally and > 500 nm axially^1, 2^. Since the size of many biological structures fall below this diffraction limit, researchers have developed a collection of super-resolution microscopy techniques to overcome this optical barrier, achieving nanometer-scale resolution ^3-7^. Although these super-resolution techniques are powerful, they often require expensive equipment, specialized techniques and reagents, and extensive post-imaging processing ^8^. A recently developed method, termed Expansion Microscopy (ExM), offers an economical alternative for enhancing the resolution of light microscopy by embedding biological samples in a polyacrylamide gel network and physically expanding the gel-sample entity via hydro-osmotic expansion, effectively improving the resolution by the expansion factor of the hydrogel-sample entity ^8-14^. ExM has proven to be versatile in enabling both two-dimensional and three-dimensional imaging of a range of biological specimens, including cellular components^15, 16, 11, 14, 17^, tissues ^18^, and even entire organisms ^19^.

ExM protocols have been optimized to address the various needs of the research community ^12, 14, 16, 17, 20, 21^. For example, the original ExM produced a 4-fold expansion, however more recently optimized methods provide 10∼20-fold expansion by modifying hydrogel components or iterative expansion^10, 14, 20, 22, 23^. A common technical challenge in ExM imaging is obtaining accurate 3D stack images because the samples become significantly larger and thicker after expansion. Since samples are embedded into soft and fragile hydrogels, ExM samples often exhibit a physical drift during 3D image acquisition^13, 24-27^. ExM sample drift arises from the fact that the expanded hydrogel is ∼99% water and the nature of its composition causes it to slide, even when securely mounted in a tightly sealed imaging chamber. Since ExM sample drift is a recurring problem, many experimental solutions have been proposed to address this issue including removing excess water around the gel (which can shrink the gel), poly-lysine coating the cover glass (which has not produced reliable results), agarose mounting (which can shrink the expanded gel), and even super glue^24, 25, 27^. Despite the proposed solutions, none have emerged as a reliable solution and, unfortunately, ExM sample drift remains relatively unaddressed. Here, we developed 3D-Aligner to computationally address the issue of ExM sample drift by performing background feature detection and matching between adjacent z-planes for image registration throughout the z-stack. 3D-Aligner determines sample drift, corrects images for sample drift, and outputs the drift-corrected 3D images for downstream analysis. User-defined parameters and options are seamlessly integrated into the workflow, ensuring ideal and optimal conditions for drift correction. We demonstrate the performance of the 3D-Aligner drift correction algorithm on 3D images acquired through our recently optimized 3D-ExM methods^14^ (see **Methods**). These optimized 3D-ExM methods provide approximately 4-fold or 12-fold 3D-isotropic expansion, showcasing the robustness of 3D-Aligner across different expansion samples and image registration methodologies.

## Results

### ExM samples often experience physical drift during 3D image acquisition

Imaging hydrogel-embedded biological specimens can be technically challenging due to the physical drift of hydrogels (**Extended Data Fig. 1a**). The magnitude and direction of drift in ExM images vary randomly across different samples, ranging from negligible to hundred microns throughout a z-stack. An example of drift in ExM images (12-fold expansion) of a human retinal pigment epithelial (RPE1) cell nucleus, stained for DNA and centromeres (CENP-C), is shown in **Fig. 1a-b**. In this example, the sample drifted over 70 μm in the y-direction and only a few microns in x-direction, resulting in significant distortion in the reconstructed image of the centromere structure in the yz-view (**Fig. 1b**). Given that precise quantification of biological structures critically relies on accurate 3D image reconstruction, it is important to correct the image distortion caused by sample drift.

**Figure 1.**
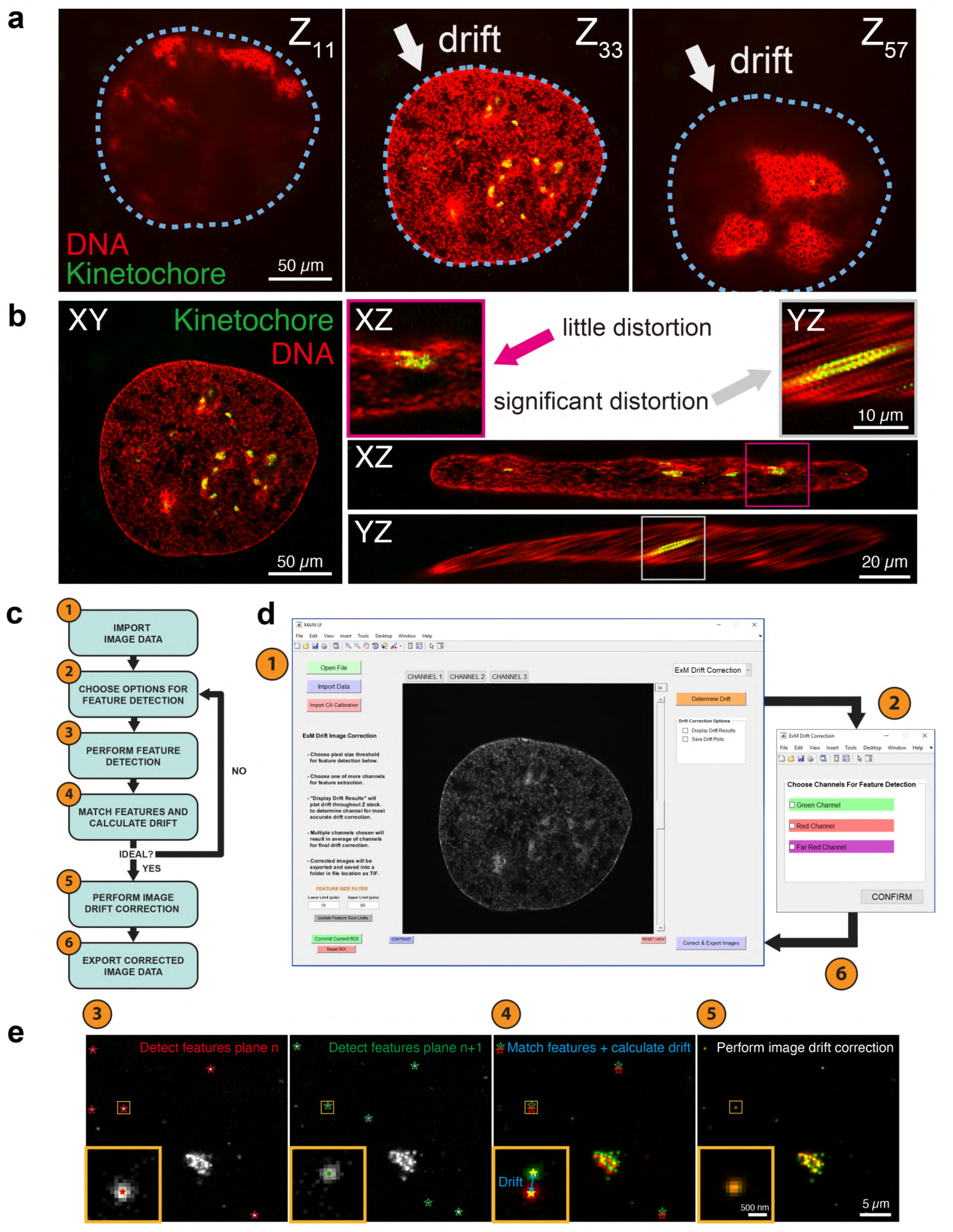
The workflow of 3D-Aligner. **(a)** Representative 12-fold expanded ExM images of an RPE1 nucleus labeled DNA and kinetochores experiencing sample drift. White arrows denote the direction of the drift and blue dotted line represents the nuclear boundary. **(b)** Images of xy, xz, and yz views of the drifted RPE1 nuclei in (a) showing the impact of sample drift on 3D image reconstruction. **(c)** Diagram showing the 3D-Aligner workflow. (1) Import images, (2) choose options for feature detection, (3) perform feature detection, (4) match features and calculate drift, (5) perform image drift correction, and (6) export corrected image data. **(d)** Snapshots of the 3D-Aligner interface and workflow. **(e)** Representative images showing feature detection, feature matching, drift calculation, and image drift correction between adjacent z planes. Stars mark detected features from background signals.

### 3D-Aligner determines the drift with nanometer precision

To correct for image distortion caused by drift, we developed a MATLAB-based package called 3D-Aligner (**Fig. 1c-e**). 3D-Aligner is integrated into 3D-Speckler, a software we introduced for microscope calibration and nanoscale measurements^2^. The workflow of 3D-Aligner involves 6 simple steps: (1) import images, (2) choose options for feature detection, (3) perform feature detection, (4) match features and calculate drift, (5) perform drift correction, and (6) export corrected image data (**Fig. 1c**). The user interface is simple, allowing the entire process to be completed with minimal clicks (**Fig. 1d**). Users can customize options for feature detection, such as selecting feature sizes and expected drift magnitude, to accommodate the range of expansion factors in different ExM methods. Feature detection is exclusively performed on isolated fluorescent speckles in the background of the image, such as those resulting from non-specific antibody staining, rather than on larger biological structures, to prevent image correction artifacts (**Extended Data Fig. 1b**). Features detected in a specific z-plane are aligned with corresponding features in adjacent z-planes. Subsequently, the drift is calculated to enable precise 3D image correction (**Fig. 1e**).

Drift correction is performed from the calculated cumulative drift at each z-plane. **Fig. 2** presents an example pre- and post-3D-Aligner corrected image of a 12-fold expanded RPE1 interphase centromere. In this example, the centromeres drifted over 16 μm along the x-direction and 4 μm along the y-direction across the z-stack. Images showcasing centromeres before (labeled red) and after (green) correction at the 48, 82, and 132 z-planes are presented. These results demonstrate that 3D-Aligner effectively measures sample drift between adjacent z-planes across a 3D image stack.

**Figure 2.**
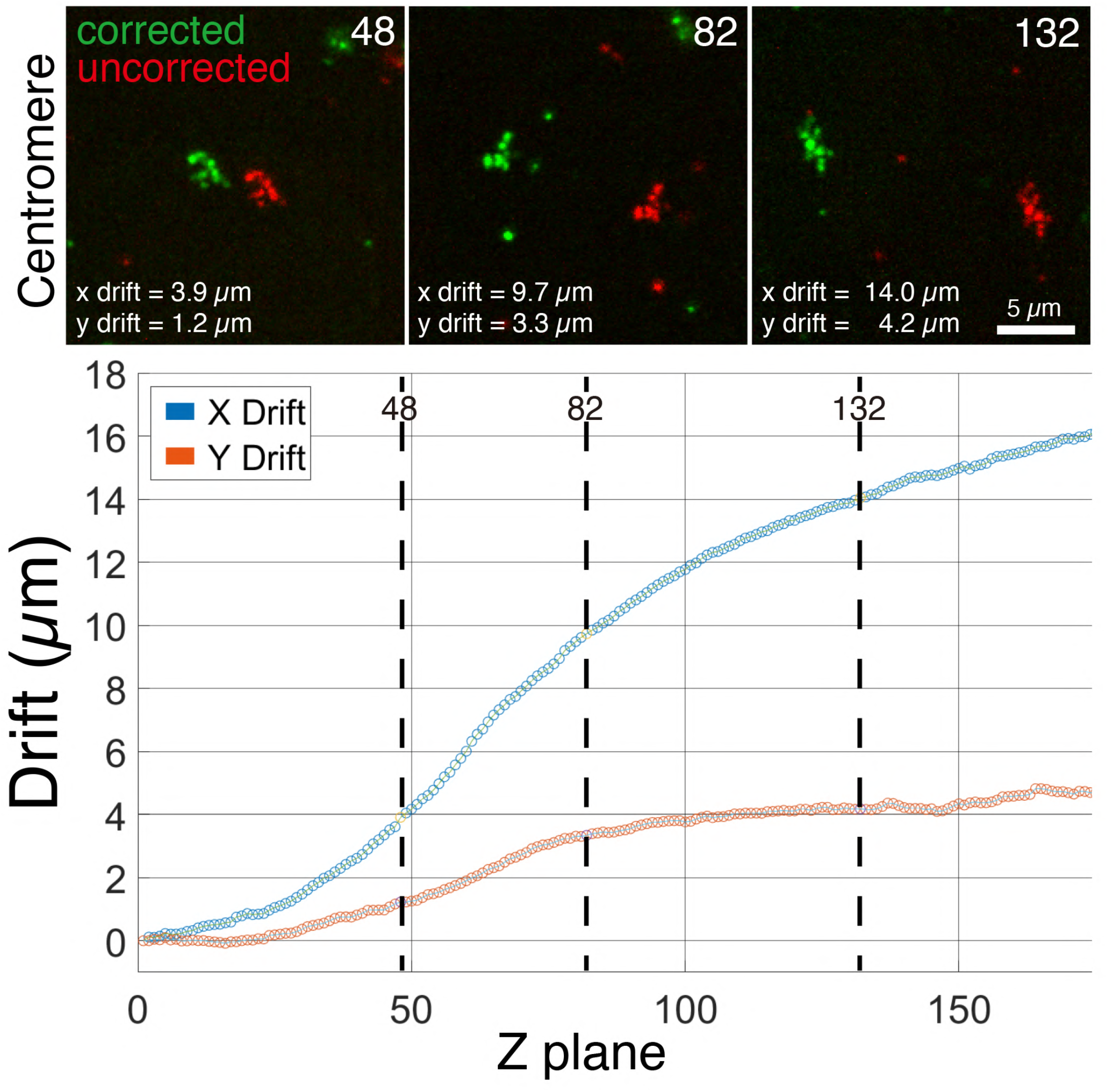
3D-Aligner determines the magnitude of drift in ExM images with high precision. Representative pre- and post-corrected 12-fold expanded ExM images of centromeres labeled with CENP-C in RPE1 cells (top) and plot showing the calculated cumulative drift in the x and y-axes (bottom).

We next investigated the consistency of calculated drift values across different colored channels of independently labeled targets. We hypothesized that the drift during ExM image acquisition would be comparable across different color channels within the same imaging plane when similar exposure times are used. To test this hypothesis, we used 3D-Aligner to measure the drift in 4-fold and 12-fold ExM images labeled with different centromere antibodies (against CENP-C and CENP-B) or a combination of centromere and microtubule antibodies (against ACA and Tubulin). Our analysis revealed that the differences in the average drift between channels, under all above staining conditions, were consistently less than 8 nm in both the x- and y-directions (**Extended Data Fig. 2a-b**). These results support the concept that drift correction values obtained from a single channel can be effectively applied to other color channels acquired under comparable exposure time for 3D image drift correction. Collectively, they validate that 3D-Aligner determines sample drift with high precision.

Proper feature detection for drift determination can depend on the dyes and antibodies used. For example, labeled microtubules distribute a wider range along the z-axis, allowing for the detection of features throughout the entire z-stack (**Extended Data Fig. 2b, left**). On the other hand, labeled nuclei was confined to a narrower range along the z-axis compared to cytoplasmic components (such as microtubules), making feature detection difficult beyond this axial range (**Extended Data Fig. 2b, middle**). Feature detection in the centromere channel enabled drift determination over a broader axial range compared to the DNA channel (**Extended Data Fig. 2b, right**). This is likely due to the antibody generating non-specific background over a wider range than the DNA dye. When multiple labeled targets occupy different z-ranges and feature detection issues are present in multiple channels, 3D-Aligner offers an option to calculate the average drift at each z-plane across several, user-defined channels (**Extended Data Fig. 2c-d**). Average drift correction can suppress errors in calculated drift due to feature detection issues (**Extended Data Fig. 2d**). These results demonstrate the capabilities of 3D-Aligner in correcting drift in ExM images, utilizing drift correction values obtained from either a single channel or the average of multiple channels.

### 3D-Aligner effectively corrects sample drift and accurately reconstructs 3D structures

We further evaluated the advanced performance of 3D-Aligner for drift correction and 3D image reconstruction in both 4-fold and 12-fold ExM images. In the 12-fold ExM image, RPE1 cells were stained with a centromere antibody and a DNA dye (**Fig. 3a, left**). The sample exhibited a drift of ∼0.6 μm per z-step along the y-direction and ∼0.03 μm along the x-direction, as determined from the centromere channel (**Fig. 3b, left**). This resulted in significant distortion of both centromere and nuclear structures along the yz-axis compared to the xz-axis (**Fig. 3a, left)**. The image after drift correction by 3D-Aligner, using values obtained from the centromere channel, are shown in **Fig. 3a, right**. Remarkably, the 2D and 3D architectures of both the nucleus and centromeres were significantly improved following drift correction. To quantitatively assess the effectiveness of drift correction, we re-applied 3D-Aligner to the corrected 3D image stack, confirming that the drift was substantially suppressed in the corrected image stack (∼0.003 μm / z-step along both x and y-axes) (**Fig. 3a, right, Fig. 3b, right, and Extended Data Fig. 3a**). An additional example of a 12-fold ExM image, before and after correction by 3D-Aligner, is shown in **Extended Data Fig. 3b**. This image before drift correction exhibited a drift of approximately 0.1 μm per z-step along the x-direction and 1.3 μm per z-step along the y-direction. After applying the drift correction, these values were significantly reduced to ∼0.003 μm and 0.025 μm per z-step along the x- and y-directions, respectively.

**Figure 3.**
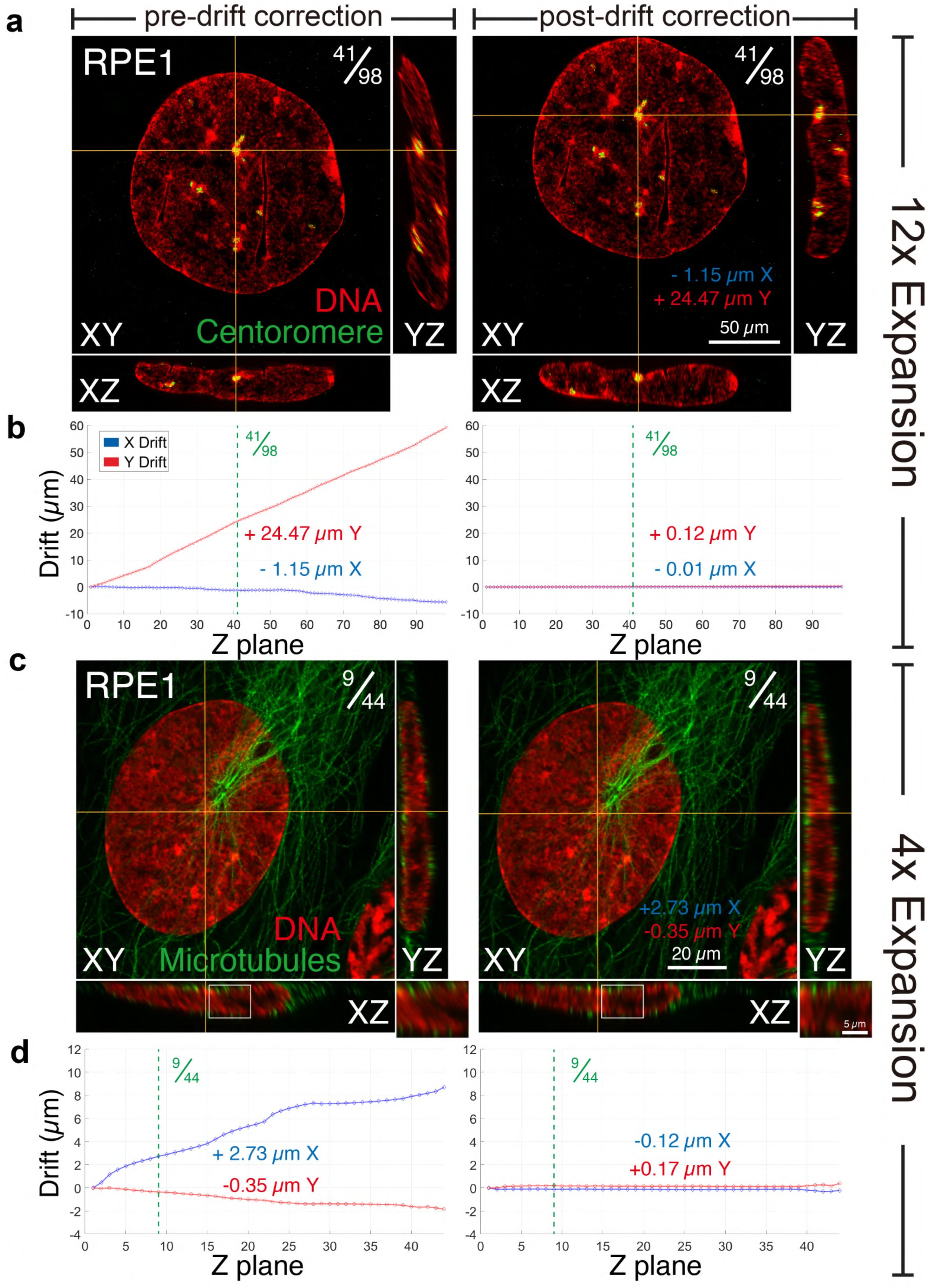
3D-Aligner effectively corrects drift in both 4-fold and 12-fold expanded ExM images. **(a)** Representative pre- (left) and post-drift corrected (right) 12-fold ExM images of RPE1 cells labeled DNA and centromeres (CENP-B). **(b)** Plots showing calculated cumulative drift in the x and y-axes across the z stack pre- (left) and post- (right) drift correction. **(c)** Representative pre- (left) and post-drift corrected (right) 4-fold ExM images of RPE1 cells labeled DNA and microtubules. Zoomed images show distortion pre- and post-drift correction. **(d)** Plots showing calculated cumulative drift in the x and y dimensions across the z stack pre- (left) and post- (right) drift correction.

We further validate the effectiveness of 3D-Aligner’s drift correction on a 4-fold expanded ExM image, where RPE1 cells were stained with both a tubulin antibody and a DNA dye (**Fig. 3c**). Unlike the previous example of a 12-fold ExM image, this sample exhibited non-uniform drift in the x-direction. Impressively, 3D-Aligner successfully suppressed the drift in this scenario, demonstrating performance consistent with the previous example (**Fig. 3c-d, and Extended Data Fig. 3a**). **Extended Data Movies 1 and 2** showcase the 3D stack images of before and after 3D-Aligner correction of both the 4-fold and 12-fold expanded ExM images.

We demonstrated significant differences in the size and volume quantifications of labeled nuclear and centromere structures before and after 3D-Aligner correction. A pre-correction 12-fold expanded ExM image of a nucleus and centromere displayed significant distortion along the drifted direction, which was significantly improved post-correction (**Fig. 4a**). The thickness of the nucleus, depending on its location, could result in a two-fold underestimation (**Fig. 4a, i and i’**) or a slight overestimation (**Fig. 4b, ii and ii’**) in the pre-correction image. Additionally, the centromere length was significantly overestimated before correction (**Fig. 4b, iii and iii’**). To further investigate the impact of drift correction on the quantification of biological structures, we measured the length of centromeres in multiple 12-fold expanded cells. Consistent with measurements in **Fig. 4b**, the longest centromere length was significantly overestimated due to drift (**Fig. 4c**). Beyond the 2D structure, 3D-Aligner improved visualization and volumetric quantification of the 3D structures of centromeres and nuclei (**Fig. 4d**). These results demonstrate that physical drift in ExM can cause significant errors in quantification, and that 3D-Aligner effectively suppresses these errors by correcting sample drift. In conclusion, 3D-Aligner accurately determines and corrects for sample drift in ExM, producing drift-corrected images suitable for precise 2D and 3D reconstitution and downstream quantitative analyses.

**Figure 4.**
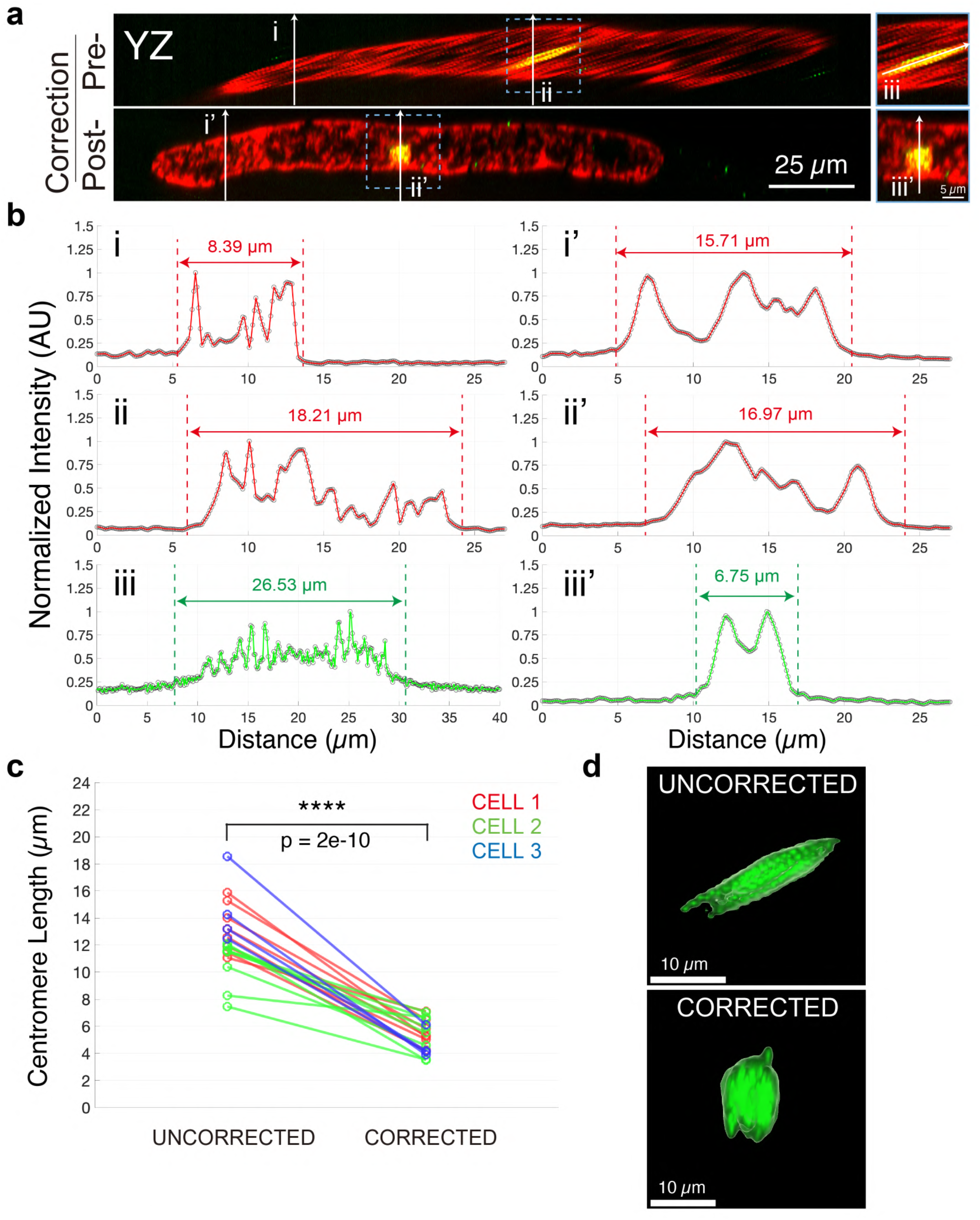
Effects of drift correction on quantification of biological structures. **(a)** Representative pre- (top) and post-drift corrected (bottom)12-fold ExM images of RPE1 cell labelled DNA and centromeres. Blue boxes show zoomed images highlighting effects of drift on a centromere. **(b)** Intensity profiles of line scans drawn in (a) to highlight quantification of structures pre- (i, ii, iii) and post- (i’, ii’, iii’) drift correction. Dotted lines highlight boundaries of either the nucleus (i, i’, ii, ii’) or centromere (iii, iii’). **(c)** Plot showing centromere lengths as measured in (b) of the same centromeres pre- (left) and post- (right) drift correction. N = 20 kinetochores across 3 cells. **(d)** Representative images of surfaces fit over the same kinetochores pre- (top) and post- (bottom) drift correction using IMARIS.

## Discussion

ExM is increasingly gaining popularity in cell biology research, with protocols continuously evolving in diverse directions to meet the varied needs of researchers. ExM enhances resolution by physically expanding specimens in 3D, necessitating a large field of view (FOV) with an extensive z-range for imaging. This physical expansion significantly reduces fluorescence signals, leading to extended image acquisition times. These prolonged imaging durations and large z-ranges further increase the risk of distortions from physical drift, making accurate post-image processing for drift correction essential. In this study, we introduce 3D-Aligner, an innovative and user-friendly image correction tool that computationally addresses sample drift in ExM, ensuring accurate 3D image reconstruction and downstream quantification. We underscore the importance of correcting for sample drift by demonstrating substantial errors in the visualization and measurement of centromere and nucleus size and volume between pre- and post-corrected images (**Fig. 4**).

3D-Aligner employs a nearest neighbor-based matching algorithm^28^ to align feature sets between adjacent z-planes, instead of relying on a pattern-matching algorithm^29, 30^. While 3D-Aligner effectively corrects for drift in most of ExM images tested, the nearest neighbor algorithm can struggle with feature matching between z-planes if there is a sudden increase in drift magnitude, though such occurrences are rare during 3D image acquisition. Furthermore, feature matching can be erroneous in densely populated feature sets where the average distance between features within the same plane is less than or equal to the drift magnitude between adjacent z-planes. To minimize these errors, 3D-Aligner allows the user to adjust the nearest neighbor threshold value based on drift magnitude and the size range of detected features according to feature density. Further improvements in feature matching can potentially be achieved by implementation of a robust pattern-matching algorithm. Feature detection may be limited in channels where staining lacks punctate background spots, which is often the case in channels imaging highly specific dyes. For the drift correction in these images, we recommend including at least one antibody-stained channel that spans a broader z-dimension for feature detection and subsequent image correction in other color channels.

3D-Aligner effectively corrected for drift in both 4-fold and 12-fold expanded ExM images with tissue culture cells and embryos tissue samples, showcasing its robust capability in drift correction for broad ExM imaging. Furthermore, 3D-Aligner software holds significant potential for future applications, including tracking the movement of fluorescent particles and correcting structural distortions in non-ExM imaging conditions. 3D-Aligner can determine the position of features with an accuracy of up to 2 nm since it uses 3D-Speckler functions for determination of positions ^2^. This capability enables the precise tracking of a moving fluorescent particle with nanometer accuracy in live imaging. An example of particle tracking using 3D-Aligner is illustrated in **Extended Data Fig. 4**, demonstrating its ability to detect bead movements even in slightly out-of-focus images. Overall, 3D-Aligner provides a powerful tool for distortion correction in ExM super-resolution images, ensuring precise visualization and facilitating accurate downstream quantification.

## Acknowledgement

We would like to thank Drs. Nathan Claxton, Hiroshi Nishida, Yoshitaka Sekizawa, Yokogawa Electrical Corporation, Nikon Japan, and Nikon USA for critical equipment and technical support. Part of this work is supported by Wisconsin Partnership Program, the University of Wisconsin-Madison Office of the Vice Chancellor for Research with funding from the Wisconsin Alumni Research Foundation (WARF), WARF Big Data Challenge Grant, start-up funding from University of Wisconsin-Madison SMPH, UW Carbone Cancer Center, and McArdle Laboratory for Cancer Research, and NIH grant R35GM147525 (to A.S.).

## Author contributions

J.L. developed the code under the guidance of A.S.. J.L., D.G., and X.Q. performed experiments and data analysis. A.S. designed and supervised the entire project, initiated work, contributed key ideas, designed experiments. J.L., D.G. and A.S. drafted the manuscript. All authors revised and contributed to the manuscript.

## Methods

### 3D-Aligner Code Availability

3D-Aligner is developed in MATLAB and is built into 3D-Speckler as a package. 3D-Aligner source code and detailed instruction are available at GitHub: https://github.com/suzukilabmcardle/3D-Aligner This software utilizes the Bio-Formats (OME) software tool for opening and reading over 140 different microscopy image file formats ^31^. However, the size of Bio-Formats exceeds GitHub’s file limit. The Google Drive link provided below contains the source code, the Bio-Formats file, and a sample image for review: https://drive.google.com/drive/folders/15_nXhPmW60pvuuSI_lxOB-VsXEWtwGwD?usp=share_link

### Feature Detection and Matching

Edges are detected via Laplacian of Gaussian and morphologically closed with a 2×2 structuring element. Detected objects are filtered by referencing the intensity values in the original image and filled. Features are filtered by size based on user-defined size ranges (default 10 < size < 60). We recommend 10 < size < 60 pixels for 12-fold expanded ExM images and 10 < size < 45 pixels for 4-fold expanded ExM images. Features are matched between adjacent z planes via a nearest neighbor algorithm and the drift between each plane is calculated as the median of the differences in positions between matched points in adjacent planes.

### Intensity Profile Generation

Intensity profiles were generated via line scans drawn in Nikon Elements software. Line scans were draws across kinetochores connecting the lowest point of the kinetochore in the lowest z plane to the highest point of the kinetochore in the highest z plane.

### Cell Culture

RPE1 cells, which were originally obtained from ATCC, were grown in Dulbecco’s modified Eagle’s medium (DMEM; Gibco) or DMEM-F12 medium (Gibco) supplemented with 10% FBS (Sigma), 100U/ml penicillin, and 100 mg/ml streptomycin (Gibco) at 37 °C in a humid atmosphere with 5% CO_2_.

### ExM images

Detailed protocols of both 4x and 12x 3D-ExM were described in Roshan et al^14^. Briefly, the 4x 3D-ExM hydrogel was prepared using acrylamide and N, N-methylenebisacrylamide (MBAA), which were used in the original ExM method^9^. The 12x 3D-ExM hydrogel was consists of N, N-dimethylacrylamide (DMAA) and sodium acrylate (SA), as used in previous study^12^. RPE1 cells were fixed with 3% PFA in PHEM buffer (120 mM Pipes, 50 mM HEPES (Sigma), 20 mM EGTA, 4 mM magnesium acetate [pH 7.0]) at 37 °C without pre-permeabilization. Cells were then rinsed with 0.5% NP40 and incubated in 2% BSA (Sigma, A4287)) and Boiled Donkey Serum (Jackson ImmunoResearch, 017-000-121). The following antibodies and dyes were used: guinea pig anti-CENP-C (MBL, PD030), mouse CENP-B (santacruz, sc-22788), mouse anti-tubulin (Sigma, DM1A, 05-829), minimum cross-react donkey anti-guinea pig (Jackson ImmunoResearch, 706-546-148), mouse (Jackson ImmunoResearch,715-086-150), DAPI (Sigma, D9542).

### Imaging

For confocal image acquisition, a high-resolution Nikon Ti-2 inverted microscope equipped with a Yokogawa SoRa CSU-W1 spinning disc confocal and a Hamamatsu Flash V2 CMOS camera was used. A 60x water immersion objective (NA/1.3) was used for ExM imaging. Most of 3D-stack images were obtained sequentially with 400 nm z-steps using Nikon Elements software.

### Image analysis

Visualization of images and Figure 4b-c analysis were performed Nikon Elements software and Imaris (Andor).

### Statistics

All statistical tests for significance between two conditions were performed using two-tailed t-tests.

